# Diet-induced hyperplastic expansion in subcutaneous adipose tissue and protection against adipose progenitor exhaustion in female mice are lost with ovariectomy

**DOI:** 10.1101/2024.09.05.611480

**Authors:** Taylor B. Scheidl, Jessica L. Wager, Jennifer A. Thompson

## Abstract

**Background:** The protection of females against cardiometabolic disease is in part attributable to a tendency for fat accumulation in subcutaneous depots, which promote lipid homeostasis by serving as a metabolic sink. At menopause this protection is lost, and body fat distribution resembles the male-like pattern of visceral adiposity. Adipose progenitor cells (APCs) can be recruited to support adipose expansion in the setting of obesity. Sex differences in diet-induced APC responses may in part explain sexual dimorphism in risk for obesity-associated insulin resistance; however, the role of sex and estrogen in governing APC function remains unclear.

**Methods:** Four groups of C57BL/6 mice were assessed: intact males vs. females, and sham vs. ovariectomized (ovx) with or without 17β-estradiol (E2). Adipogenesis was stimulated by rosiglitazone (rosi), while obesity was induced by high fat/fructose diet (HHFD). Flow cytometry quantified the total number of APCs and identify committed preadipocytes by the loss of CD24 expression. Body composition was measured by NMR, while adipose function assessed by measuring circulating adipokines and free fatty acids and lipolysis in adipose explants.

**Results:** Despite greater accumulation of fat mass in response to rosi, females were protected against the depletion in subcutaneous APCs and preadipocytes that was observed in rosi-treated males.

Similar to intact males, APC and preadipocytes in subcutaneous depots of ovx females were reduced after rosi treatment. The protection of obese females against the development of insulin resistance and adipose dysfunction was lost with ovx, and E2 re-supplementation rescued HFFD- induced APC exhaustion. Exposure to HFFD after discontinuation of rosi exacerbated glucose intolerance in males only.

**Conclusions:** Estrogen-mediated hyperplastic expansion in subcutaneous depots permits renewal of the APC pool and preservation of adipose function.

**PLAIN ENGLISH SUMMARY:** Despite well-established sex differences in the risk for type 2 diabetes that vary across the lifespan, very little is known regarding sex-specific mechanisms in its pathophysiology. In the setting of obesity, stem cells resident in fat tissue can be recruited for the generation of new fat cells, an important mechanism that maintains metabolic health. It is thought that a reduced availability or dysfunction in fat-residing stem cells is an important pathophysiological event that triggers the onset of obesity-associated type 2 diabetes. Herein, we aimed to determine how sex and estrogen influence stem cell availability and function. Our data show that the ability of fat- residing stem cells to respond to an obesogenic environment is greater in females in an estrogen-dependent manner. Estrogen-dependent stem cell responses to an obesogenic environment may contribute to the protection of females against obesity-induced type 2 diabetes and loss of this protection after menopause.

**HIGHLIGHTS:** Sexual dimorphism in activation of adipogenesis by rosiglitazone is mediated by estrogen.

Exhaustion of the APC pool occurs in subcutaneous depots of male mice, while estrogen mediates protection of females against APC exhaustion.

Preservation of subcutaneous adipose expansion capacity due to renewal of the progenitor pool may contribute to protection of females against obesity-associated insulin resistance.

## BACKGROUND

The Atlas Report by the World Obesity Federation estimated that in 2020, 38% of the global population was overweight or obese, and predicted this number to rise to ∼50% over the next decade. Across all levels of genetic risk, obesity is the most significant driver of type 2 diabetes (T2DM) (1,2). While in some populations the prevalence of obesity is greater in females, sexual dimorphism in patterns of body fat distribution that arise at puberty endow females with a relative protection against obesity-associated cardiometabolic disease. Males have a higher tendency to accumulate white adipose tissue (WAT) in the abdominal region and in visceral depots, whereas females favour WAT storage in the gluteo-femoral region and in subcutaneous depots (3,4). High waist circumference and waist-to-hip ratio are surrogate markers of central obesity and better predictors of T2DM than body mass index (BMI) (3,5).

Thus, the distribution pattern of WAT characteristic of pre-menopausal females likely contributes to their lower risk for cardiometabolic disease at a given BMI (3,4). This female advantage wanes during menopause (6–8), at which time WAT distribution shifts to a more ‘male like’ pattern (4). Further, studies of menopausal human females and ovo-hysterectomized non-human primates demonstrate that the use of estrogen-based hormone replacement therapy restores preferential fat storage in subcutaneous depots and ameliorates metabolic dysfunction associated with low estrogen (9–11). Together, these findings suggest an important role for sex steroids in regulating body fat distribution and cardiometabolic risk in females. Despite these well-established sex differences in body fat distribution patterns and metabolic disease risk, female animals have been systematically excluded from fundamental investigations into WAT function in both health and disease (12). For this reason, current understanding of the pathophysiology of obesity is largely built on the male model and there remains a paucity of knowledge surrounding sex-specific mechanisms.

It is now recognized that the risk for obesity-induced cardiometabolic disease is not only related to the distribution of WAT, but also the ability of WAT to carry out its central role in promoting lipid homeostasis. Remodeling and expansion of WAT is an adaptive process that preserves lipid homeostasis in the face of an obesogenic environment and can occur by hypertrophy of existing adipocytes and *de novo* generation of adipocytes *via* recruitment of a resident population of adipose progenitor cells (APCs). The ’adipose tissue expandability hypothesis’ posits that obesity-associated disease is precipitated by failure of the APC pool to accommodate further expansion of WAT (13). Traditional thought held that adipogenesis occurred primarily in subcutaneous depots, based on the fact that APCs isolated from subcutaneous depots differentiate readily *in vitro* whereas visceral APCs are less adipogenic in adherent culture (14). These findings conformed to the adipose tissue expandability hypothesis as subcutaneous depots play a protective role by acting as the body’s primary metabolic sink.

More recently, transgenic models for inducible adipocyte labelling that allow assessment of adipogenesis *in vivo* revealed that high fat diet-induced *de novo* adipogenesis occurs predominately in visceral depots (15,16). These landmark studies were performed in male mice and there is at least one study showing that the lack of adipogenesis in subcutaneous depots is male-specific (17), although this finding has been recently challenged by a study reporting diet-induced APC proliferation in both subcutaneous and visceral depots in male mice (18).

Ambiguity remains with respect to the role of APCs in the pathophysiology of obesity and in the sexual dimorphism of body fat distribution patterns and risk for cardiometabolic disease. Herein, we set out to determine if the contribution of APCs to subcutaneous adipose remodeling is influenced by sex and estrogen. Our findings demonstrate that males are vulnerable to APC exhaustion and that APC exhaustion compromises metabolic health in the setting of chronic obesity. Using an ovx model, we show that in females, estrogen mediates diet-induced APC responses to pharmacological activation of adipogenesis and protects females against exhaustion of the APC pool in subcutaneous depots. Thus, this study suggests that the recruitment of tissue-resident APCs to support adaptive subcutaneous adipose remodeling is a female-specific mechanism that preserves metabolic homeostasis, while loss of APC plasticity after menopause contributes to heightened risk for cardiometabolic disease.

## METHODS

### Animals

All experiments involving animals were performed at the University of Calgary in Alberta, Canada and were approved by the University of Calgary Animal Care Committee and conducted in accordance with guidelines by the Canadian Council on Animal Care Ethics.

Reporting of animal procedures align with the ARRIVE guidelines. Animals were housed on a 12/12-hour light-dark cycle at 22.5°C and 35% humidity. Prior to 7 weeks of age, female C57BL/6J mice underwent ovariectomy (ovx) or sham surgery (sham), performed by Jackson Laboratories. Intact male and female C57BL/6J mice were bred in house. At ∼ 9 weeks of age, mice were fed high fat/high fructose diet (HFFD) *ad libitum* [45% kcal from fat, 35% kcal from fructose; Research Diets Inc. (D08040105I)] for 16 weeks to induce obesity. Control mice were fed a low-fat control diet [10% kcal from fat; Research Diets Inc. (D12450Ki)]. A subset of ovx mice were administered a physiologically relevant dose of estrogen (0.05ug/mouse in sesame oil) by subcutaneous injection every 4 days, to mimic the estrus cycle. Whole body composition was measured by time domain nuclear resonance (TD-NMR; Bruker, LF-90II). Food intake was determined following a 6h starvation period by refeeding individually housed mice with 100g of HFFD and measuring intake after 2, 6, and 18h (overnight).

In a subset of mice, adipogenesis was stimulated pharmacologically by oral administration of the PPARᵧ agonist, rosiglitazone (15mg/kg/day) (Rosi; TCI Chemicals, R0106) or a vehicle control consisting of 1% methylcellulose (Veh; Sigma Aldrich, M0512) in 50% sweetened condensed milk, for 8 weeks. Following rosi or vehicle treatment, animals were either euthanized, HFFD feeding was initiated after discontinuation of rosi. Animals were euthanized by decapitation following induction with 5% isoflurane anaesthetic with 1L/min O2. Whole blood was collected and centrifuged for 10 minutes at 2000 x g to isolate serum.

### Isolation of APCs

Inguinal subcutaneous WAT (sWAT) and gonadal visceral WAT (vWAT) were collected into HEPES + 2% BSA (Millipore Sigma, A2153) with 1mg/mL collagenase (Worthington Biochemical, LS004196) and minced by sterile surgical scissors, followed by digestion at 37°C on a tube rotator for 45 minutes. Digested tissue was then passed through a 100um filter, and the stromal vascular fraction (SVF) isolated by centrifugation following lysis of erythrocytes. SVF was either processed for downstream flow cytometry or resuspended in preadipocyte growth media (Cell Applications, 811-500) and passed through a 70um cell strainer prior to seeding in a T25 flask for *in vitro* experiments.

### Flow Cytometric Quantification of *in vivo* APC Quantity and Distribution

Freshly isolated SVF was resuspended in 1mL of PBS + 5% FBS and passed through a 70um filter to remove clumps. Cells were counted using the Countess II Automated Cell Counter (Thermofisher) and resuspended at a concentration of 0.5 x 10^6^ cells/mL. Cells were first stained with viability dye (Biotium, 32003) for 30 mins, followed by antibody staining. For quantification of APCs, cells were stained for 1h with anti-CD45 and anti-CD31 for exclusion of leukocytes and endothelial cells (Lin^-^); anti-CD34 and anti-PDGRFα to identify the general population of APCs; and anti-CD24 for differentiating uncommitted (CD24+) from uncommitted APCs (CD24-), as described by Rodeheffer *et al* (38). After staining, cells were washed by centrifugation and fixed in 1% PFA for 15 minutes. Samples were analyzed using the Cytek Aurora flow cytometer and data analyzed using FlowJo V10.0. Singlets were defined on the basis of forward (FSC) and side scatter (SSC) and gates determined by single color, unstained, and fluorescent minus one controls (FMO). Antibody concentrations and vendors can be found in table 1.

**Table 1.**
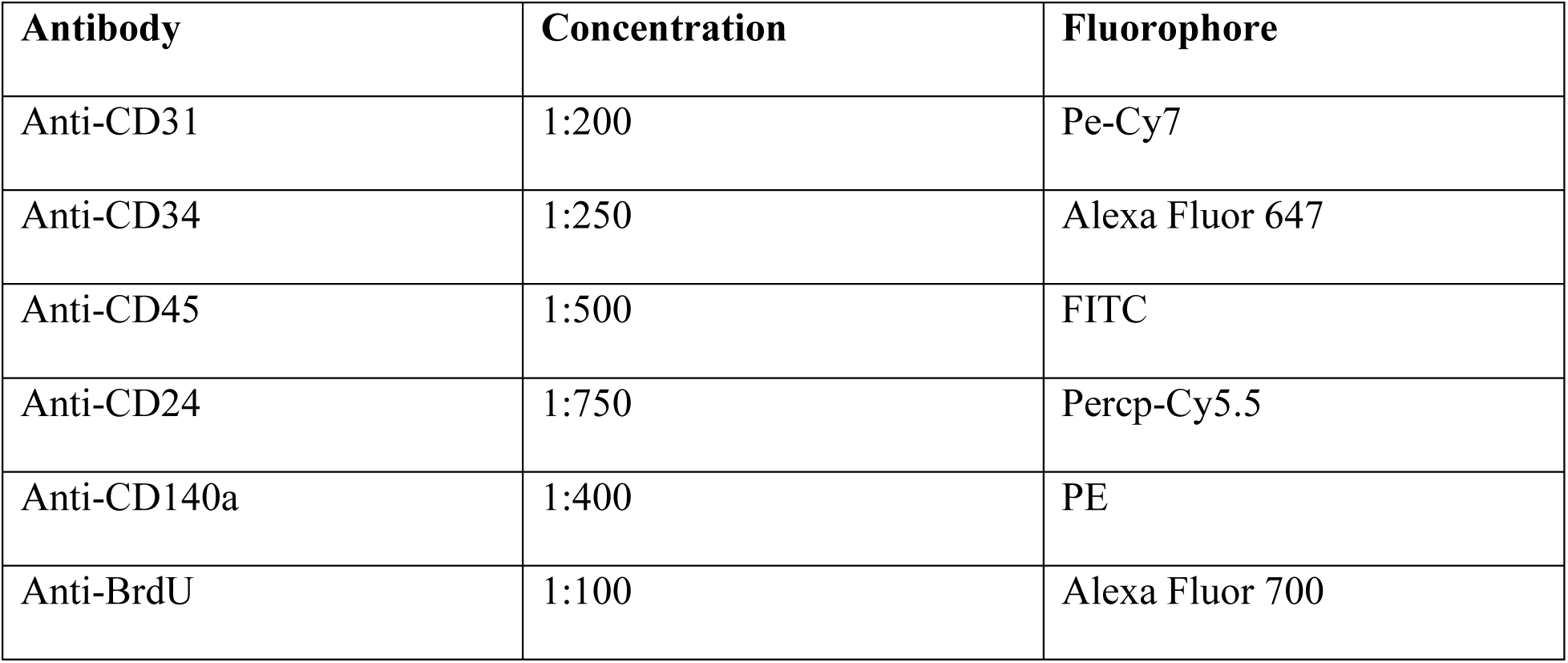
Antibodies for Flow Cytometry.

**Table 2.**
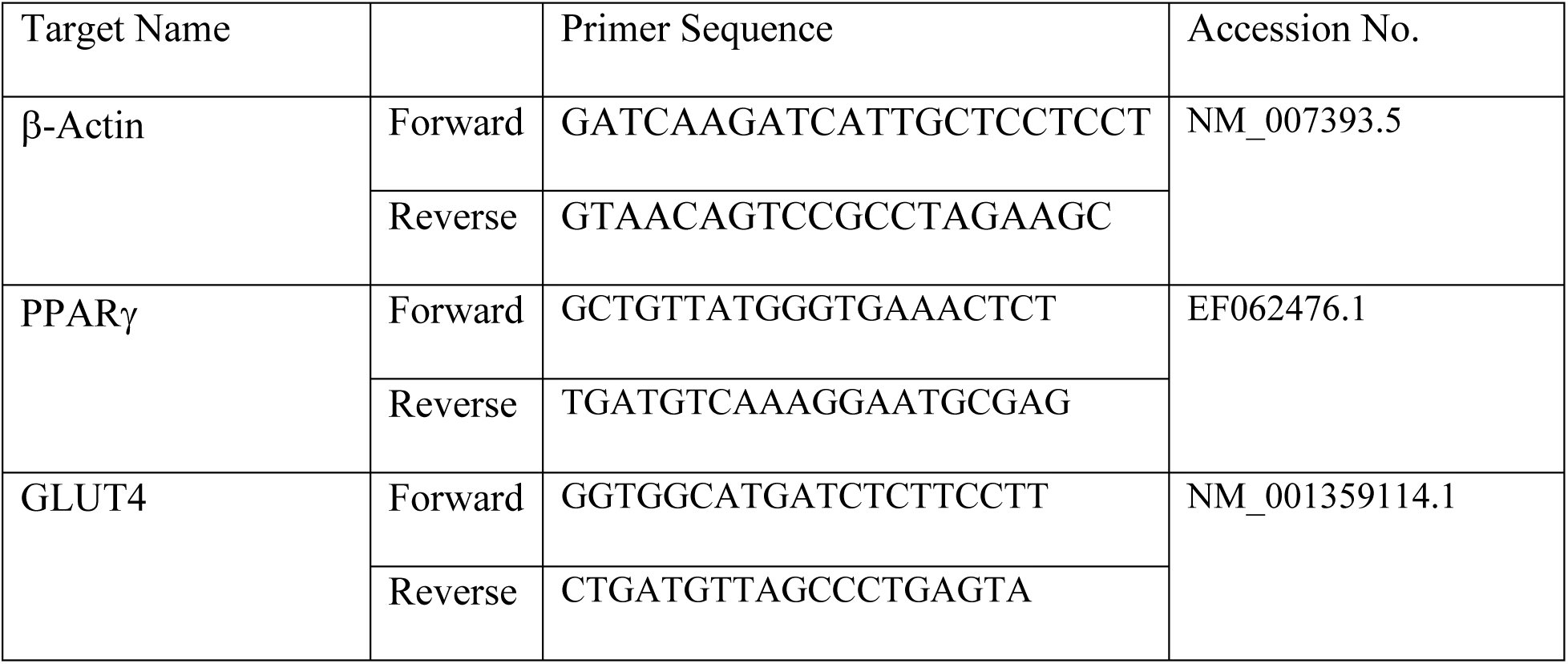
Primer Sequences and Accession Numbers.

### Whole-Body Glucose Tolerance & Serum Markers of Adiposopathy

Whole body glucose tolerance was assessed by intraperitoneal glucose tolerance test (IPGTT). Briefly, mice were starved in fresh cages for 6 hours and administered glucose (2g/kg) by intraperitoneal injection. Glucose was measured from tail blood at baseline and 5-, 10-, 15-, 30-, 60-, and 120-minutes post-injection using the AlphaTrak 2 glucometer. The circulating levels of free fatty acids (FFA) (Abcam, ab65341) and insulin (Cayman Chemical, 589501) were quantified in the serum of fasted animals following the manufacturer’s instructions. WAT insulin resistance (ADIPO-IR) was calculated by multiplying FFA x insulin (39). Adiponectin (Thermofisher, KMP0041) was similarly quantified in the serum of mice starved 16h and re-fed for 1 h.

### Adipocyte Size

Fixed sWAT and vWAT were embedded in paraffin blocks and sectioned by the University of Calgary Faculty of Veterinary Medicine Diagnostic Services Unit. Sections were stained with hematoxylin and eosin (H&E) to assess adipocyte cell size. Images were captured using the Nikon Eclipse Ts2 microscope. The ImageJ Fiji plugin Adiposoft was used to measure adipocyte size, with 4 images per animal evaluated.

### *Ex Vivo* Lipolysis

After starving mice for 16h followed by re-feeding, sWAT and gonadal vWAT was collected and small (70-90mg) pieces were aseptically dissected and placed in a 24-well tissue-culture treated plate (Greiner Bio-One, 662160), immersed in M199 (Thermofisher, 11150067) + 1% FBS (Cytiva, SH30396.03) and allowed to equilibrate overnight in a 37°C 5% CO2 incubator. The following day, media was removed, and tissue washed twice with PBS before adding 500uL of KRBH media (KRBH + 2% fatty acid free BSA [Millipore Sigma, 126609]). After 2h, media was collected to serve as a baseline and 10uM isoproterenol (Sigma Aldrich, 420355) was added to the KRBH media to induce β-adrenergic stimulated lipolysis. After 2h, media was collected and stored at -80°C. Wet tissue weight was recorded for data normalization. Glycerol release was quantified in the media samples using the Abcam Free Glycerol Assay Kit (ab65337) per the manufacturer’s instructions.

### Statistical analyses

Given the impact of surgery itself, single variable comparisons were made between intact male and intact female or between ovx and sham females using a student’s t-test. Two-way ANOVA with Sidak’s multiple comparisons test were used for analysis of time-dependent parameters or to assess the effects of treatment and sex or depot. Data are shown as mean ± SEM, and comparisons are considered significant when p<0.05. All statistical analyses were performed using Prism 10.

## RESULTS

### Rosiglitazone-induced WAT expansion is sexually dimorphic and mediated by estrogen in females

To directly measure *in vivo* adipogenic capacity, we treated mice with rosiglitazone, a potent pharmacologic inducer of adipogenesis (19,20) (Fig. 1A, B). Intact rosi-treated females gained ∼2-fold more whole-body fat mass over the 8 weeks of rosi treatment compared to vehicle-treated females (Fig. 1C). In contrast, whole body fat mass did not appreciably increase in response to pharmacological stimulation of adipogenesis by rosi in males (Fig. 1D). Loss of estrogen in females abolished the rosi-stimulated gains in body fat mass, as shown in sham vs. ovx (Fig. 1E). Comparing intact mice administered vehicle, a higher ratio of sWAT-to-vWAT in females vs. males was attributable to higher vWAT in males (Fig. 1F – H).

**Figure 1.**
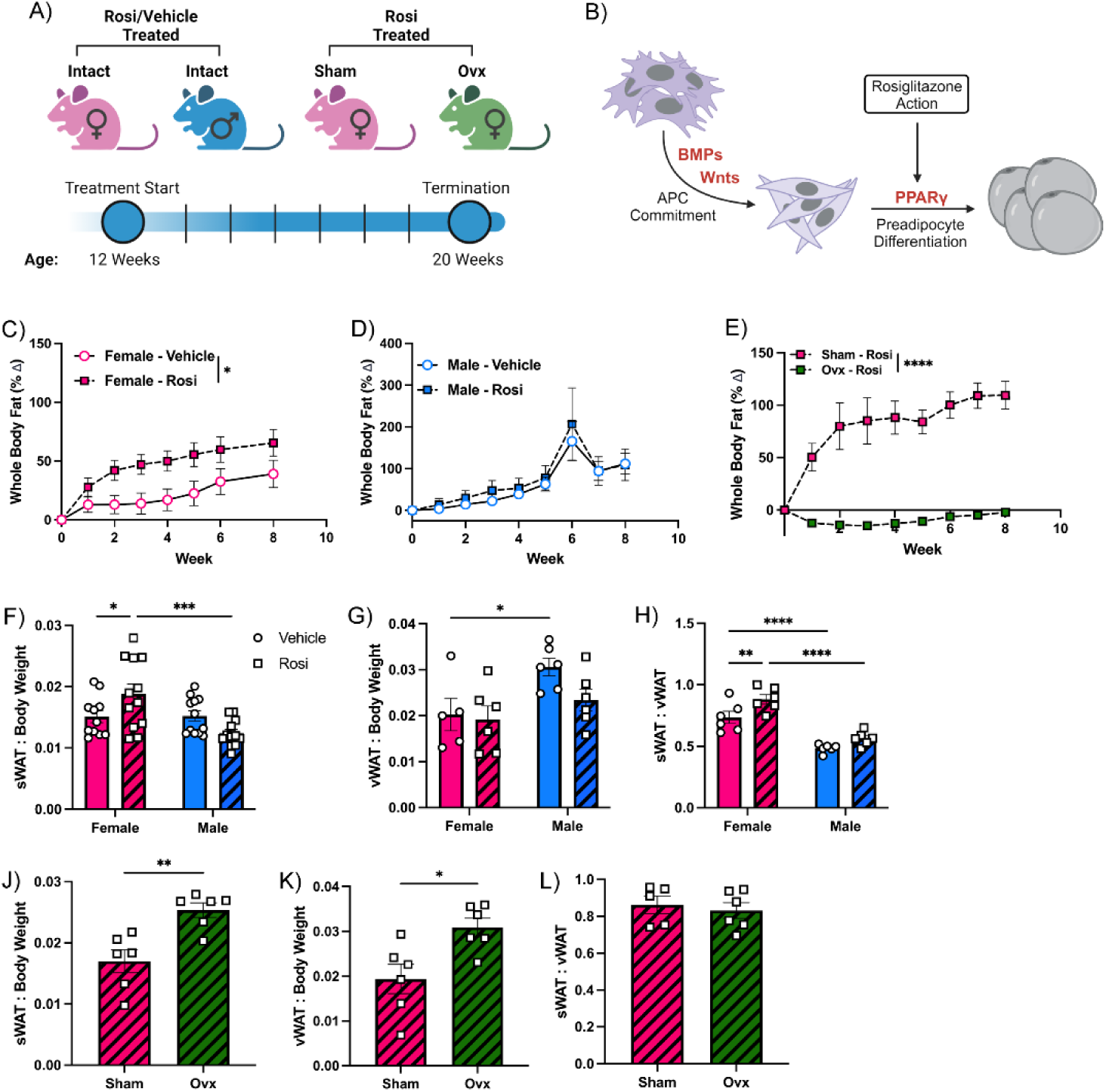
Estrogen is permissive of pharmacologically induced whole-body fat accumulation. A) Male and female mice were treated with rosiglitazone or vehicle for 8 weeks; ovx and sham mice were treated with rosi for 8 weeks. Body composition was measured weekly using TD-NMR. B) Mechanism of rosiglitazone action on APCs. Whole-body fat accumulation over 8 weeks vehicle or rosi treatment in C) female, D) male, and E) ovariectomized and sham operated mice, measured *via* TD-NMR. F) sWAT and G) vWAT weight relative to body weight in male and female vehicle or rosi treated mice. H) sWAT to vWAT ratio in male and female vehicle or rosi treated mice. J) sWAT and K) vWAT weight relative to body weight in rosi treated ovx and sham mice. L) sWAT to vWAT ratio for rosi treated ovx and sham mice (n=5-6). *p<0.05, **p<0.01, ****p<0.0001.

Rosi-induced whole body fat gain in females was driven primarily by increases in sWAT, while vWAT mass remained unchanged (Fig 1. F – H). Depot weight was unaffected by rosi treatment in intact males. The depot differences observed in males vs. females were not present in comparing sham and ovx mice, however, this is likely attributable to the impact of ovx on body fat mice that may mask the impact of rosi (Fig. 1J-L).

### Estrogen mediates female-specific protection against rosiglitazone-induced APC exhaustion

Interesting findings published by the Graff group revealed that prolonged rosiglitazone treatment led to an exhaustion in the progenitor pool and reduced adipogenic potential of APCs (19), although neither sex nor depot was reported. Data herein reveal that in the sWAT of rosi-treated males, the total number of APCs (Lin-: CD34+; PDGRFα+) are depleted compared to vehicle-treated males (Fig. 2D). Uncommitted APCs (CD24^+^) made up a greater proportion of the total APC pool in rosi-treated males, as reflected by a higher number of CD24^+^ APCs and lower number of committed preadipocytes (CD24^-^) (Fig. 2E, F). Given that rosi is a PPARγ agonist that stimulates the terminal differentiation step of adipogenesis (21), these data imply that pre-existing preadipocytes in male sWAT differentiated in response to rosiglitazone, but were not replenished by the pool of proliferating APCs. Under control conditions, female sWAT contained a lower total number of APCs compared to males, but a greater proportion of female APCs were identified as committed preadipocytes (Fig. 2D-F). Female APC populations were unaffected by rosi treatment (Fig. 2D-F) despite significant body fat gain. Female-specific protection against rosi-induced APC exhaustion was lost in ovx females, which displayed an enrichment in uncommitted CD24^+^ APCs and decrease in committed CD24^-^ preadipocytes in sWAT and vWAT (Fig. 2G-I), similar to intact males. These data suggest that estrogen is protective against rosiglitazone-induced exhaustion of the preadipocyte pool.

**Figure 2.**
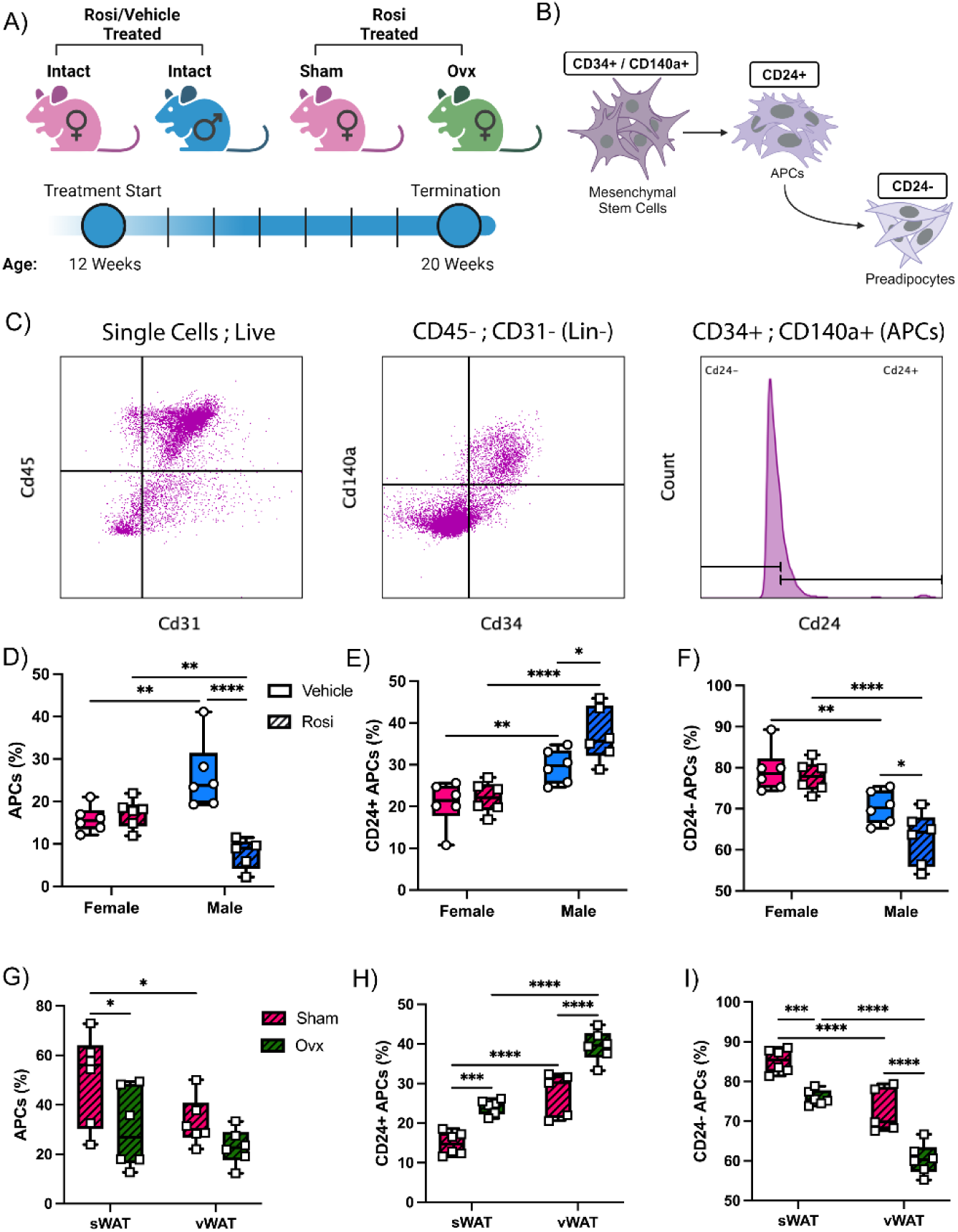
Estrogen protects against exhaustion of the adipose progenitor pool following chronic pharmacologic stimulation of adipogenesis. A) Male, female, ovx-, and sham-operated mice were given vehicle or rosi for 8 weeks. B) Schematic of APC populations quantified by flow cytometry. C) Gating paradigm for quantification of adipose progenitor cells and CD24+/- progenitors. Lin^-^ cells (CD45-/CD31-) are gated from single, live cells. CD34+/CD140a+ APCs are sub-gated from Lin- population. CD24+ and CD24- APCs are identified from double positive APC population. D) APCs in sWAT as a percentage of total live cells in male/female rosi or vehicle treated mice. E) Number of CD24+ and F) CD24- APCs in sWAT as a percentage of APCs in male/female rosi or vehicle treated mice. G) Number of adipose progenitor cells in sWAT and vWAT as a percentage of total live cells in rosi treated ovx and sham mice. H) Number of CD24+ and I) CD24- cells in sWAT and vWAT as a percentage of total adipose progenitor cells in rosi treated ovx and sham mice. (n=5-6). *p<0.05, **p<0.01, ***p<0.001, ****p<0.0001.

### Estrogen administration rescues diet induced APC exhaustion

Given that estrogen appears to protect against exhaustion of the APC pool by prolonged pharmacological stimulation of adipogenesis, we measured APC populations after 16 weeks of CD or HFFD. Females on HFFD had a significantly greater proportion of total APCs than those on CD (Fig. 3B), but diet had no effect on the APC population in males, suggesting a female-specific enrichment of APCs in response to the HFFD. Committed APCs made up a greater proportion of the total APC population in HFFD- and CD-fed females vs. males (Fig. 3D). HFFD feeding significantly increased the number of committed APCs in females when compared to those fed a CD (Fig. 3D). To determine whether this female protective effect was due to estrogen, we administered exogenous E2 to ovx females over the course of 16 weeks HFFD. E2 administration blunted body fat accumulation in response to HFFD, and minimized whole body fat mass (Fig. 3F, G). Ovx females had fewer APCs than sham females on both the CD and HFFD. E2 administration completely rescued the ovx-induced reduction in total APCs (Fig. 3H). Sham and ovx+E2 females had fewer CD24+ and greater proportion of CD24- APCs relative to ovx females, suggesting a greater propensity for adipogenesis in sham and estrogen replete females (Fig. 3I,J).

**Figure 3.**
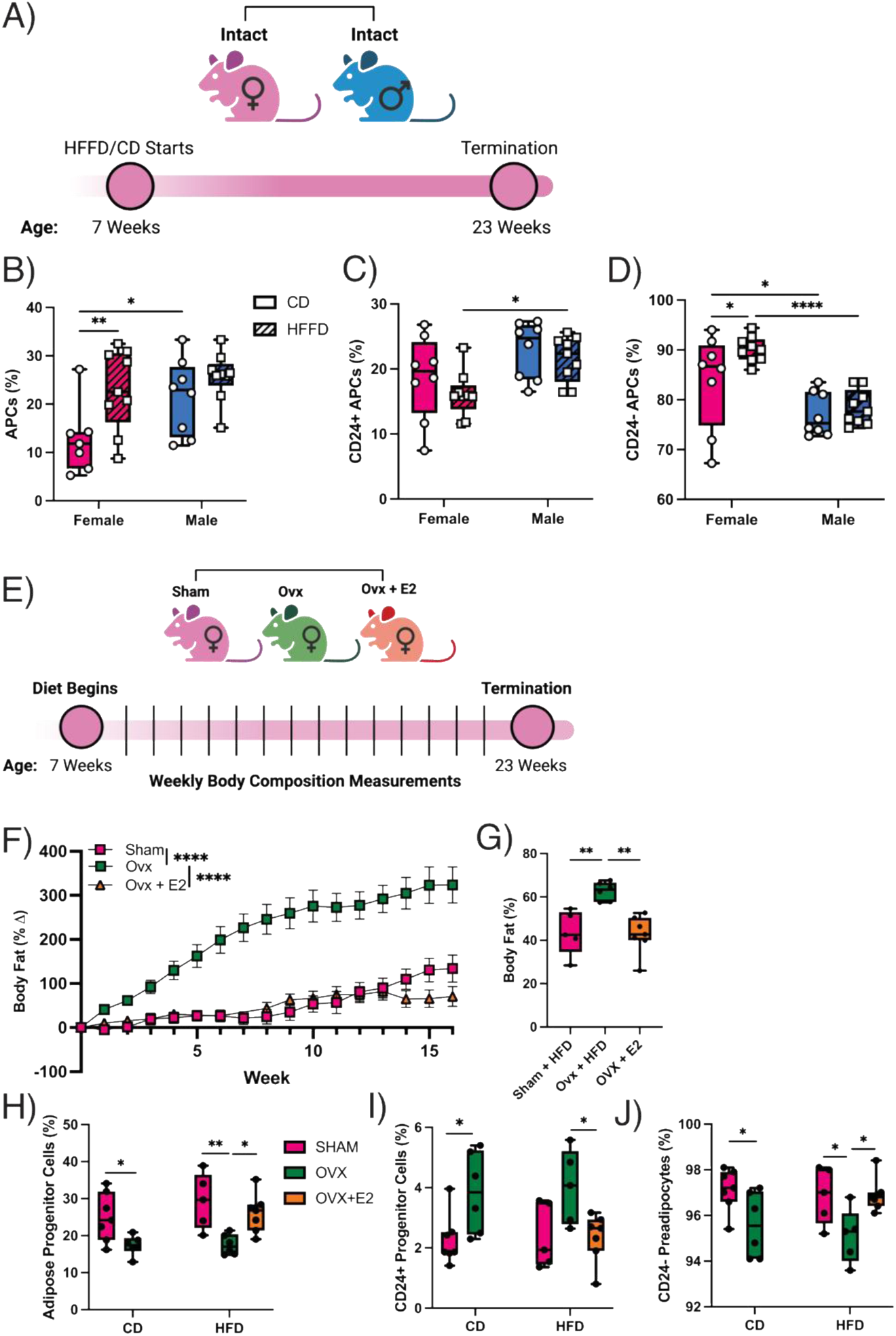
Estrogen is permissive of HFFD-induced APC self-renewal in sWAT. A) Male and female mice were fed CD or HFFD for 16 weeks. B) APCs in sWAT of CD- or HFFD-fed male and female mice, as a percentage of total live cells. Flow cytometric quantification of C) CD24+ and D) CD24- APCs as a percentage of total APCs (n=8-10). E) Sham and ovx females with or without E2 were fed CD or HFFD for 16 weeks. F) Whole body fat accumulation over the course of the HFFD. G) Total body fat at the end of the HFFD. H) Total, I) CD24+, and J) CD24- sWAT APCs. (n=5-6) *p<0.05, **p<0.01, ***p<0.001, ****p<0.0001.

### Estrogen mediates diet-induced obesity independent of food intake

It has been previously demonstrated that male mice exhibit hyperinsulinemia, impairment in insulin-mediated suppression of lipolysis, and lipid spillover after 16 weeks of HFFD, whereas females were protected against diet-induced metabolic dysfunction (22). Here, we sought to verify if this female-specific protection was mediated by estrogen. At baseline, there were no differences in body fat between intact males and females (Supp. Fig 1A). Intact males gained more whole-body fat mass over the 16 weeks of HFFD compared to females (Fig. 4B - D).

**Figure 4.**
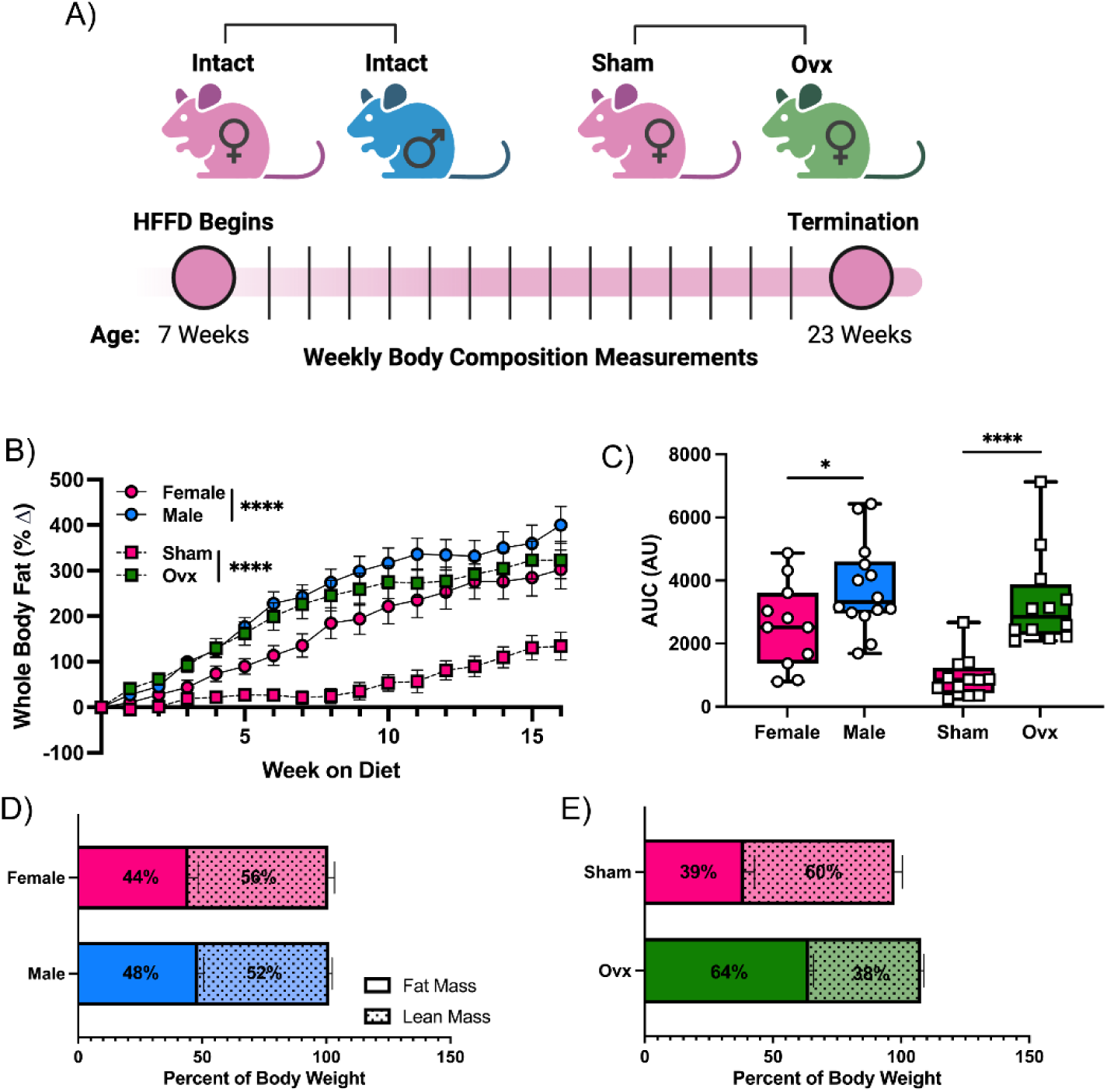
Estrogen blunts HFFD-induced body fat accumulation. A) HFFD was given for 16 weeks, and body composition measured weekly using TD-NMR. B) Percent change in whole-body fat mass from the initiation of HFFD. C) Area under the curve analysis of body fat accumulation. Lean and fat mass breakdown for D) intact males and females, and E) ovx and sham females. (n= 11-14). *p<0.05, ****p<0.0001.

Similarly, there were no baseline differences in whole-body fat between ovx and sham females, but over the course of 16 weeks HFFD, ovx females gained significantly more whole-body fat mass compared to sham-operated females (Fig. 4B, C, E), Diet-induced body fat gain was markedly less in sham-operated females compared to intact females, suggesting an impact of surgery on body composition (Fig. 4B, C). At the end of the diet period, male mice had a mean of 48 ± 2.1% body fat mass, while intact females had an average of 44 ± 3.9% body fat mass (Fig. 4D). Ovx and sham mice showed a similar pattern, with ovx mice having a mean of 64 ± 1.6% body fat and sham having a mean of 39 ± 4.1%. Changes in lean mass mirrored the differences in body fat; male mice had 52 ± 1.2%, and females 44 ± 2.4%. Lean mass in ovx females was 43 ± 1.0%, while in sham females it was 60 ± 2.5% (Fig. 1E). These estrogen-mediated differences were not accounted for by food intake; while food intake was greater in males than females, sham females had greater food intake compared to ovx females (Supp. Fig. 1B).

### Estrogen is protective against the development of obesity-associated adiposopathy

Both intact male and ovx mice developed severe HFFD-induced glucose intolerance, while glucose control remained normal in intact and sham-operated females (Fig. 5B - D).

**Figure 5.**
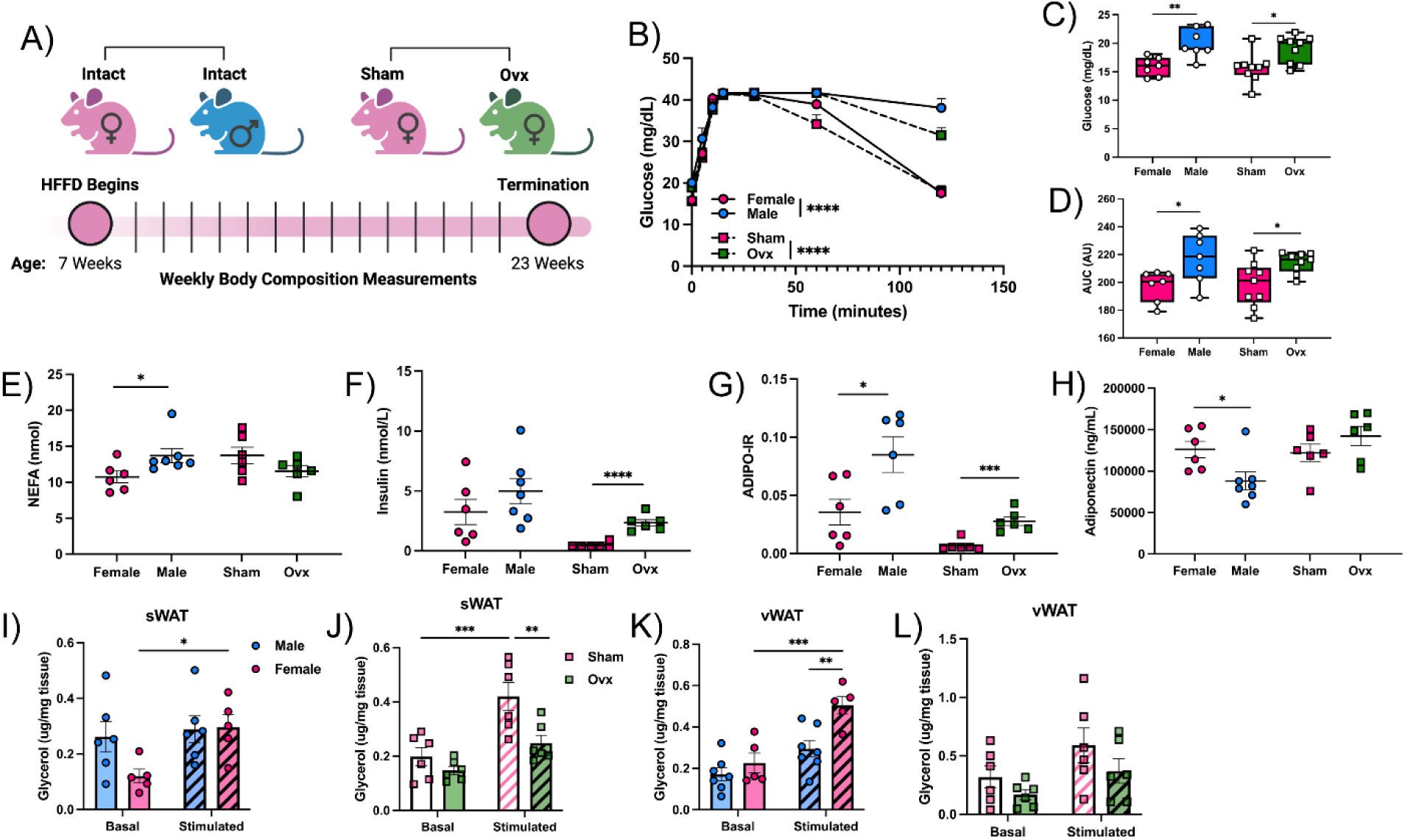
Estrogen protects against obesity-related metabolic dysfunction. A) HFFD was given for 16 weeks, tissue and serum were collected at the end of the diet. B) Blood glucose levels over the course of 2-hour glucose tolerance test. ‘HI’ readings from glucometer are shown as highest readable value, 41.7 mg/dL. C) Fasting glucose at beginning of glucose tolerance test. D) Area under the curve analysis of glucose tolerance curves. Serum levels of E) NEFA, F) insulin, G) calculated ADIPO-IR, and H) serum adiponectin. I,J) Free glycerol release from sWAT and K,L) vWAT explants at baseline and following stimulation with isoproterenol. (n=7-12). *p<0.05, **p<0.01, ***p<0.001, ****p<0.0001.

Circulating levels of NEFA were higher in HFFD-fed male mice relative to intact females but were not different between ovx and sham mice (Fig. 5E). While serum insulin levels were not statistically different between intact males and females (Fig. 5F), insulin was higher in ovx vs. sham-operated females. Postprandial insulin and NEFA were used to calculate the ADIPO-IR, a measure of the ability of WAT to supress lipolysis in response to insulin and thereby an index of WAT insulin resistance (IR). The ADIPO-IR was higher in both male and ovx compared to intact and sham females, respectively (Fig, 5G). In line with existing evidence regarding sex differences in adipokines, circulating adiponectin was higher in HFFD-fed intact females vs. males (Fig. 5H). Loss of estrogen had no impact on adiponectin levels (Fig. 5H). Relative to baseline, there was no increase in glycerol release in response to β-adrenergic stimulation in male sWAT explants (Fig. 5I), possibly due to higher basal lipolysis. In contrast, β-adrenergic stimulation significantly increased glycerol release in sWAT explants collected from intact females (Fig. 5J). Similarly, there was a significant effect of β-adrenergic stimulation on glycerol release in sWAT explants collected from sham females, which was eliminated with ovx (Fig. 5J). In vWAT explants, β-adrenergic-induced lipolysis was not observed in males, while glycerol release from vWAT explants collected from intact females was increased in response to stimulation (Fig. 5K). Ovx had no impact on basal or stimulated lipolysis in vWAT explants (Fig. 5L). Explants from HFFD-fed intact, sham, and ovx females all demonstrated a pattern of greater stimulated lipolysis in the vWAT vs. sWAT, however this difference was only statistically significant in intact and ovx females. There were no depot differences in stimulated lipolysis in explants from intact HFFD-fed males (Supp. Fig. 3A-D). In HFFD-fed intact, ovx, and sham females, adipocyte size was larger in the vWAT than sWAT (Supp. Fig. 2); while in males, adipocyte size distribution was consistent across the two depots (Supp. Fig. 2). In sWAT, intact females had a greater proportion of small adipocytes (30-40µm) when compared to males, while males had a greater proportion of larger adipocytes (60-70µm) (Supp. Fig. 4). There were no differences in ovx vs. sham groups (Fig. 4). Adipocyte size distribution was similar in vWAT across all groups (Supp. Fig. 4). These data suggest that estrogen tempers HFFD-induced adiposopathy in a depot-specific manner, possibly due to estrogen-mediated APC responses to HFFD.

#### Exhaustion of APCs in males drives vulnerability to HFFD-induced metabolic dysfunction

To determine the impact of APC exhaustion on metabolic health in the setting of obesity, we treated mice with rosi for 8 weeks, and subsequently exposed mice to HFFD for 21 weeks after discontinuation of rosi. Both male and female mice administered rosi for 8 weeks had a significantly blunted body fat accumulation in response to subsequent HFFD when compared to their vehicle treated counterparts (Fig. 6B, D). Despite limited body fat accumulation in female mice pre-treated with rosi, whole body glucose tolerance was preserved following 21 weeks of

**Figure 6.**
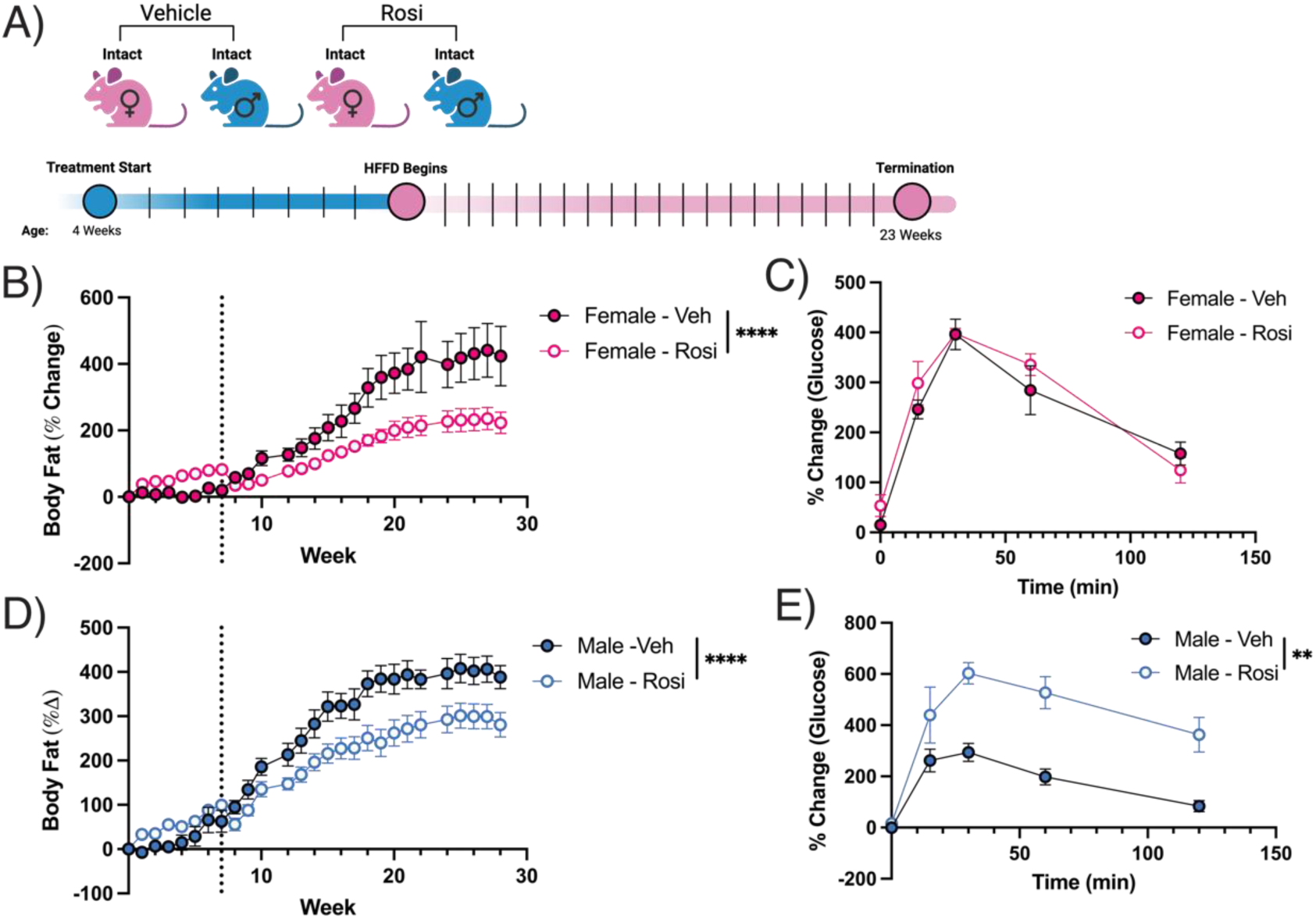
Exhaustion of APCs in males drives vulnerability to HFFD-induced metabolic dysfunction. A) rosi or vehicle were administered for 8 weeks, after which treatment was terminated and HFFD feeding began, continuing for 21 weeks. B, D) Whole body fat mass measured weekly. Dotted line indicates when rosi/vehicle treatment ended and HFFD began (n=4-6). C, D) Whole body glucose tolerance, assessed by glucose tolerance test at the end of the HFFD (n=4-6). **p<0.01, ****p<0.0001.

HFFD (Fig. 6C). Comparatively, male rosi-treated mice had significantly blunted whole body glucose tolerance after the subsequent HFFD compared to vehicle treated males (Fig. 6E).

## DISCUSSION

The goal of the present study was to determine the impact of sex and estrogen on depot-dependent APC plasticity. Our findings demonstrate that a higher capacity for WAT expansion in subcutaneous WAT depots in females is mediated by estrogen. This estrogen-mediated preservation of the APC pool permits adaptive WAT remodeling, promoting sexual dimorphism in the capacity for adipose tissue to adapt to an obesogenic environment. Thus, our findings highlight a potential mechanism contributing to the protection of females against obesity- associated cardiometabolic disease and loss of this protection after menopause (Fig. 7).

**Figure 7.**
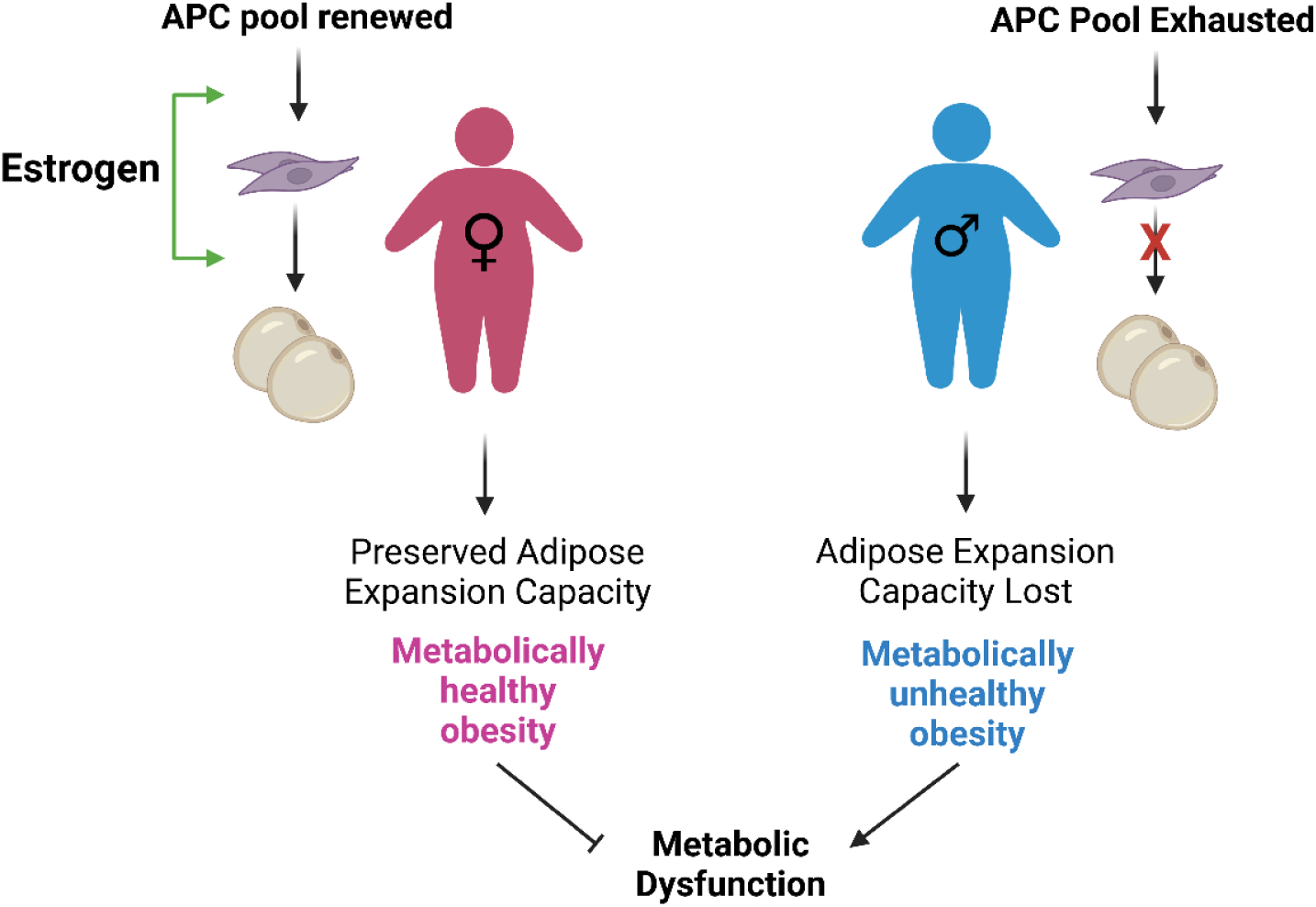
A permissive effect of estrogen on hyperplastic expansion in subcutaneous may underlie sex differences in obesity-associated cardiometabolic disease. Greater capacity for hyperplastic expansion in subcutaneous depots of female mice may be mediated by estrogen and in part contribute to the relative protection of pre-menopausal females against obesity-associated cardiometabolic disease, while males may be prone to adipose progenitor exhaustion. Impairment of hyperplastic expansion capacity due to loss of estrogen may in part underlie heightened risk for obesity-associated cardiometabolic disease after menopause. Image created in BioRender.

This study underscores the need to integrate the depot and sex-specific behaviour of APCs into the ‘adipose tissue expandability hypothesis’, a simplified model that describes the role of APCs in the pathophysiology of obesity. According to the adipose tissue expandability hypothesis, a failure in the recruitment of resident APCs for *de novo* generation of new adipocytes under conditions of high demands for lipid storage triggers the development of WAT dysfunction. Dysfunctional adipose tissue or adiposopathy characterized by chronic inflammation, insulin resistance and dysregulated lipolysis leads to spillover of lipids into the blood and deposition into ectopic organs and visceral depots. Adiposopathy is thought to distinguish metabolically unhealthy obesity from metabolically healthy obesity, the latter more common in human females while in their reproductive years (3). Although this theory is widely accepted, there exists little direct evidence supporting a causative role for APCs in the development of obesity-associated adiposopathy. The fact that APCs isolated from subcutaneous depots differentiate more readily in culture compared to visceral APCs led to the conclusion that APC differentiation under obesogenic conditions occurred predominately in subcutaneous depots. This dogma persisted until studies by the Rodeheffer and Scherer groups were published, demonstrating with transgenic techniques of inducible adipocyte labeling that adipogenic responses to an obesogenic environment occur predominately in visceral depots with negligible responses in subcutaneous depots (15,16). Likewise, Kim *et al*. used a stable isotope method of measuring adipocyte turnover and showed that the robust hyperplastic responses to an obesogenic stimulus observed in juvenile mice were attenuated in adulthood and that hyperplastic expansion was greater in visceral compared to subcutaneous depots (23), highlighting the limited plasticity of APCs in post-pubertal mice, particularly in subcutaneous depots. These landmark studies have shaped current thinking with respect to the contribution of APCs to adipose expansion in obesity, but since they did not evaluate sex as a biological variable, they are unable to determine if lack of metabolically induced hyperplastic expansion in subcutaneous depots is universal or sex specific. In a subsequent study, the Rodeheffer group compared the sexes and found that diet-induced APC proliferation and adipogenesis occurs in both subcutaneous and visceral depots in female mice and that ovariectomy abolished diet- induced APC proliferation (17). In agreement, we show herein that the increase in the total number of APCs and committed preadipocytes observed in subcutaneous depots of intact obese females was absent in obese males and that ovariectomy decreases the total number of subcutaneous APCs. Further, administration of exogenous estrogen rescues APC exhaustion in ovx females. Thus, the present study corroborates findings of Jeffrey *et al*. and suggests that adipogenic capacity of APCs is retained in subcutaneous depots of adult females, providing a potential mechanism explaining the protection of premenopausal females against obesity- associated metabolic disease.

There exists some indirect evidence linking impaired adipogenesis to the pathogenesis of obesity-associated metabolic disease. Comparing two mouse strains with different susceptibilities to obesity-induced insulin resistance, Kim *et al*. showed that insulin sensitivity after prolonged high fat diet was associated with higher capacity for sWAT adipogenesis (23). In obese human females, greater adipogenic potential of APCs isolated from subcutaneous, but not visceral depots, was predictive of lower metabolic syndrome risk (24). Two recently published studies using single cell RNA sequencing to examine the heterogeneity of APCs revealed a rearrangement in APC subpopulations in diet-induced obesity. The first study reported a lower number of immature proliferative APCs and impaired *in vitro* differentiation capacity of committed preadipocytes, in epididymal depots, but not subcutaneous depots, of male mice that were obese and glucose intolerant. This finding lends support to the idea that obesity is associated with impaired adipogenic potential and a failure to replenish the APC pool in a depot that undergoes diet-induced hyperplastic expansion. In subcutaneous depots of male mice, the absence of hyperplastic expansion may account for the lack of effect of diet-induced obesity on APC availability, as shown by the study above and data presented herein. A study by Cho *et al*. showed that immature APCs made up a greater proportion of total APCs while the proportion of preadipocytes was decreased in epididymal fat pads of male mice after 13 days of high fat diet. This shift in the balance of uncommitted and committed APCs after short-term high fat diet may be due to the diet-induced APC proliferation that occurs over the first week of high fat diet and precedes APC differentiation (15), rather than a pathophysiological change in differentiation capacity or preadipocyte availability that might be expected with the metabolically unhealthy obesity that manifests with prolonged high fat diet feeding. Impaired adipogenic capacity of APCs in the setting of severe obesity and co-morbid insulin resistance may be attributable to a phenotypic change or premature senescence in an environment of chronic low grade adipose inflammation. For example, an exhaustion in APC and preadipocyte reservoirs have been observed in subcutaneous depots of ageing mice and coincide with an emergence of a subpopulation of pro-fibrotic, anti-adipogenic APCs (25).

Interestingly, a study by Tang *et al.* demonstrated that prolonged stimulation of terminal adipocyte differentiation with rosiglitazone suppressed the differentiation capacity and renewal of tissue resident PPAR-γ expressing APCs without increasing cell death (19), although neither sex nor depot was reported. These data suggest that APC exhaustion may occur with stimulation of adipogenesis in an adipose tissue environment wherein inflammation is absent. Herein, we used the same duration and dose of rosiglitazone and show that in both intact males and estrogen-deficient ovariectomized females, prolonged stimulation of adipogenesis exhausts the total pool of APCs and decreases the proportion of committed preadipocytes in subcutaneous depots. These data suggest that, in males, existing preadipocytes may have terminally differentiated into adipocytes upon pharmacological stimulation but were not replenished by APC proliferation. Rosiglitazone-induced APC exhaustion in males resulted in exacerbated glucose intolerance with subsequent exposure to an obesogenic environment, suggesting that APC exhaustion does indeed elicit a vulnerability to diet-induced metabolic dysfunction. However, this appears to be a uniquely male phenomenon.

Intact and sham-operated females were protected against rosiglitazone-induced APC exhaustion, possibly due to greater APC plasticity in subcutaneous depots that allowed renewal of the progenitor pool. Rosiglitazone-induced body fat accumulation occurred exclusively in intact and sham-operated females, predominately in subcutaneous depots. Ovariectomy abolished rosiglitazone-induced body fat gain in females, suggesting that estrogen-mediated APC differentiation is the primary driver of fat accumulation. Rosiglitazone belongs to the class of thiazolidinediones, synthetic PPARγ agonists with insulin sensitizing effects in part attributable to stimulation of adipogenesis. In a small group of diabetic patients including both males and females, thiazolidinediones increased the ratio of subcutaneous to visceral fat area measured by MRI, although a sex-based analysis was not performed (26). Another study comparing the efficacy of anti-diabetic drugs in human males and females demonstrated that thiazolidinediones were associated with greater weight gain in obese and non-obese females compared to males, and elicited a greater improvement in blood glucose in females and in obese versus non-obese diabetics (27). Higher efficacy of thiazolidinediones in females may be attributable to greater APC plasticity in subcutaneous depots of females. Despite their insulin sensitizing effects, thiazolidinediones increase the risk for cardiovascular death (28) and for this reason were suspended from European markets. The present study and previous findings from Tang *et al*. demonstrate that prolonged treatment with thiazolidinediones exhausts the subcutaneous APC pool, particularly in men and post-menopausal women. While APCs have a nominal contribution to metabolically induced expansion of male subcutaneous adipose depots, a depleted reservoir of APCs would nonetheless fail to support normal adipocyte turnover, accelerating the progression of ageing-associated adiposopathy.

A permissive effect of estrogen on APC activation in gluteo-femoral subcutaneous depots and preferential accumulation of fat in this area provides an evolutionary advantage because abnormally low subcutaneous fat stores compromises fecundity, while high gluteo-femoral fat stores accumulated during pregnancy are later mobilized for lactation (29). Sexual dimorphism in body fat distribution patterns arise in puberty, driven by the surge in sex steroids. Puberty also coincides with a period of high APC proliferation (30) and thus may be a critical phase during which estrogen-mediated APC responses are established. Estrogen may modulate APC responses indirectly through influencing other cells within the adipose microenvironment.

In data presented herein, female mice have lower visceral adiposity compared to males and previously published data show protection against insulin resistance and dysregulated lipid homeostasis in female mice made obese by the Western diet used in the current study (22). Thus, mice model sexual dimorphism in body fat distribution patterns and risk for obesity-associated metabolic disease. In line with existing evidence in rodents, we show that females gain slightly less whole-body fat mass than males in response to prolonged high fat diet, while depletion of estrogen by ovariectomy leads female mice to accumulate fat mass to a similar degree as intact male mice. Although the magnitude of obesity in ovariectomized females may have been dampened by effect of the surgery itself, as suggested by the markedly lower body fat mass in sham-operated females compared to intact females. It is important to recognize that the impact of ovariectomy on diet-induced obesity and metabolic dysfunction is mediated by the pleiotropic actions of estrogen. Estrogen exerts significant influence over energy balance and diet-induced adipose expansion in a depot-specific manner through both central and peripheral mechanisms.

Estradiol interacts with estrogen receptors expressed in the nucleus of the solitary tract (NST) of the brain stem and nuclei of the hypothalamus to modulate potency of anorexigenic and orexigenic signals, suppress food intake and increase energy expenditure (32,33) . Our data demonstrate that sham-operated females have greater food intake than ovariectomized females, while intact males have greater food intake than intact females. It is important to note that these differences are of small magnitude (∼3% different). Published studies have suggested that elevated food intake in ovx mice is a transient event, and that over a longer period there are in fact no differences in food intake between ovx and sham animals (33,34). Thus, these small differences, while statistically different, may have arisen from the method of measurement.

Further, it is important to note that ovariectomy may exert its obesogenic effect through mechanisms that are both estrogen-dependent and independent. The rise in follicle stimulating hormone (FSH) levels that occurs with ovariectomy in rodents and human menopause may contribute to the heightened susceptibility to gains in body fat mass as well as impairments in glucose metabolism, the latter demonstrated in the study by Cheng *et al*. published in this series (35). Thus, while loss of estrogen may exert influence over WAT expansion and function *via* APC dynamics, there are both central and peripheral effects of estrogen and other hormones which likely contribute to the metabolic outcomes of ovariectomy.

At the level of the adipocyte, estrogen mediates expression of genes involved in the regulation of lipolysis and lipogenesis, the balance of which differs across the depots and between the sexes (4). Diet-induced glucose intolerance observed in intact males and ovariectomized females in the current study were both associated with a suppression of insulin-mediated lipolysis as reflected by higher ADIPO-IR. In obese males, β-adrenergic-induced lipolysis was absent in both subcutaneous and visceral adipose explants, possibly due to an increase in basal lipolysis and decrease in catecholamine-stimulated lipolysis, as exhibited by both rodent models of obesity and obese humans. β-adrenergic-stimulated lipolysis was retained in both depots of obese females but lost in only subcutaneous depots of ovariectomized female. An influence of estrogen on catecholamine-induced lipolytic responses in isolated subcutaneous adipocytes has been previously demonstrated, in agreement with our results (36). Overall, our findings show that adipose function is preserved in obese females but compromised with loss of estrogen. In agreement with studies showing that sex differences in adiponectin occur independent of obesity and insulin resistance, our data show higher serum levels of adiponectin in obese females compared to males. Circulating adiponectin, an anti-inflammatory and insulin sensitizing adipokine, was not impacted by the loss of estrogen and thus does not contribute to the insulin resistance in ovariectomized females.

A few limitations of this study are worth noting. First, while the ovx model is commonly used to study menopause, it fails to recapitulate the gradual decline in sex steroid production.

Rather, it causes an abrupt and near complete arrest of ovarian hormone production. Thus, the present study elucidates the role of estrogen in regulating APC behaviour and availability but cannot necessarily draw conclusions about human menopause. Many studies use a very limited number of markers to identify APCs (e.g., PDGRFα), likely resulting in contamination from non-adipose cell types, limiting conclusions with respect to APC quantity and information on subpopulations. Our flow cytometric panel was designed to accurately capture WAT resident APCs based on validated cell surface markers (37,38). Rodeheffer *et al.* demonstrated that PDGRFα, commonly used singularly to identify APCs, was only capable of marking ∼75% of these cells. In comparison, CD34+/PDGRFα+ cells gated from the Lin- population identified >90% of APCs (37). The Rodeheffer group showed that CD24+ APCs were able to generate functional WAT when injected into depots of lipodystrophic mice and differentiate into CD24- APCs, the latter expressing late adipogenic markers, suggesting that CD24 expression is lost in committed preadipocytes (38). However, a caveat of the use of flow cytometry or single cell RNA sequencing to draw conclusions regarding APC dynamics or abundance is that measurements are only captured at a single point in time.

## PERSPECTIVES AND SIGNIFICANCE

Despite a wealth of evidence indicating that cardiometabolic risk differs between the sexes and across reproductive life stages of females, research in the field of obesity has primarily relied on a male model of research, obscuring sex-specific pathophysiological mechanisms.

Herein, we demonstrate that estrogen acts permissively of APC responses to adipogenic stimuli, particularly in subcutaneous depots. This capacity for APC differentiation and renewal may be a key mechanism underlying the protection of females against obesity-related metabolic disorders and loss of this protection after menopause. Males are prone to APC exhaustion due to limited capacity for APC renewal in subcutaneous depots; however, since the contribution of hyperplastic expansion to subcutaneous adipose expansion in males is negligible in the setting of obesity, APC exhaustion may be more relevant to the pathogenesis of ageing-associated adiposopathy. The adipose tissue expandability hypothesis should be revised to incorporate the complexities that arise when considering the influence of sex on depot-dependent adaptive remodeling.

## DECLARATIONS

### Ethics approval and consent to participate

All procedures involving animals were approved by the University of Calgary Animal Care Committee (Protocol #: AC21-0132) and conducted in accordance with guidelines by the Canadian Council on Animal Care Ethics.

### Consent for publication

Not applicable.

### Availability of data and materials

The datasets used and analysed during this study are available from the corresponding author on reasonable request.

### Author’s contributions

TBS contributed to study design, performed animal experiments, collected and analyzed the data and wrote the manuscript. JLW contributed to animal experiments and provided editorial input for the manuscript. JAT contributed to conception of the study and experimental design, performed immunostaining of adipose sections, and contributed to writing and editing of the manuscript.

## Acknowledgements

We acknowledge the Flow Cytometry Core in the Cumming School of Medicine for support and use of the Cytek Aurora Flow Cytometer, particularly support and training from Drs. Ranjan Maity and Yiping Liu. We acknowledge Sophie Yonan for her assistance in analyzing adipose sections.

---

**Supp. Figure 1.**
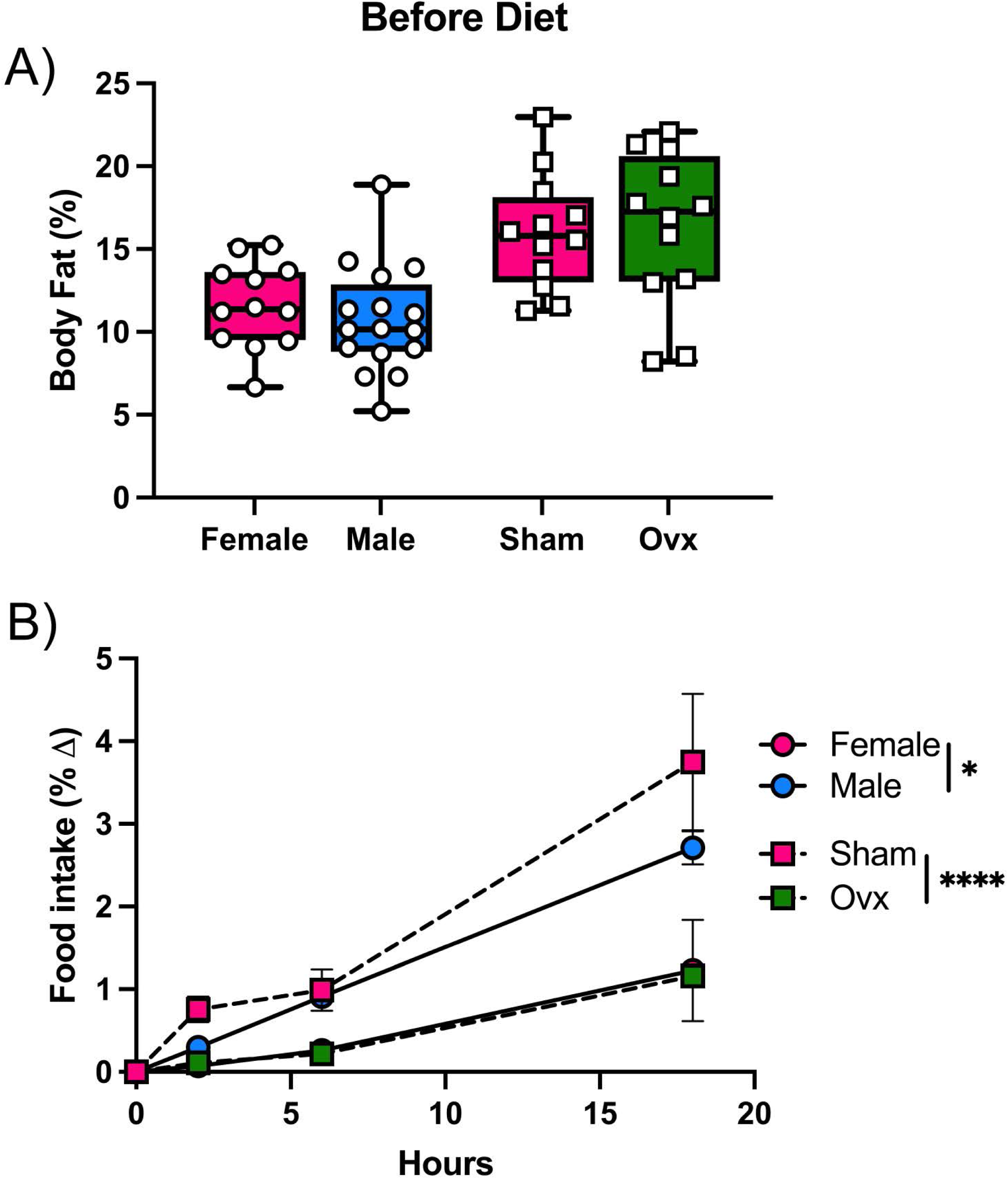
Baseline body fat and food intake for experimental model. A) baseline whole-body fat percentage of male, female, ovx, and sham mice. B) Food intake following 6h starvation period and 18-hour refeeding period in male, female, ovx, and sham mice. (n=11-14). *p<0.05, ****p<0.0001.

**Supp. Figure 2.**
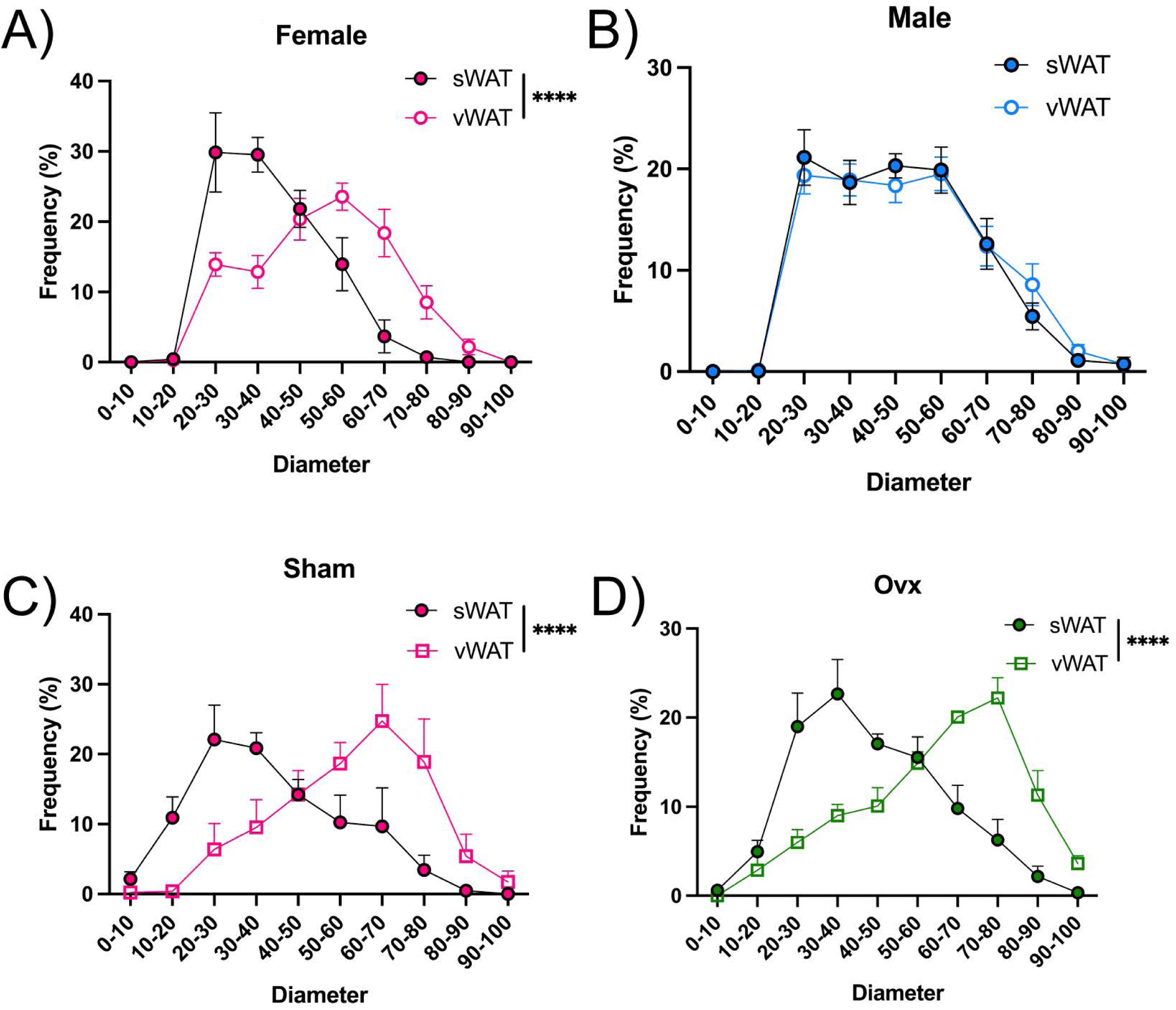
Adipocyte size is not regulated by estrogen. Adipocyte cell size comparison between sWAT and vWAT for A) female, B) male, C) sham, and D) ovx mice. (n=5-6). ****p<0.0001.

**Supp. Figure 3.**
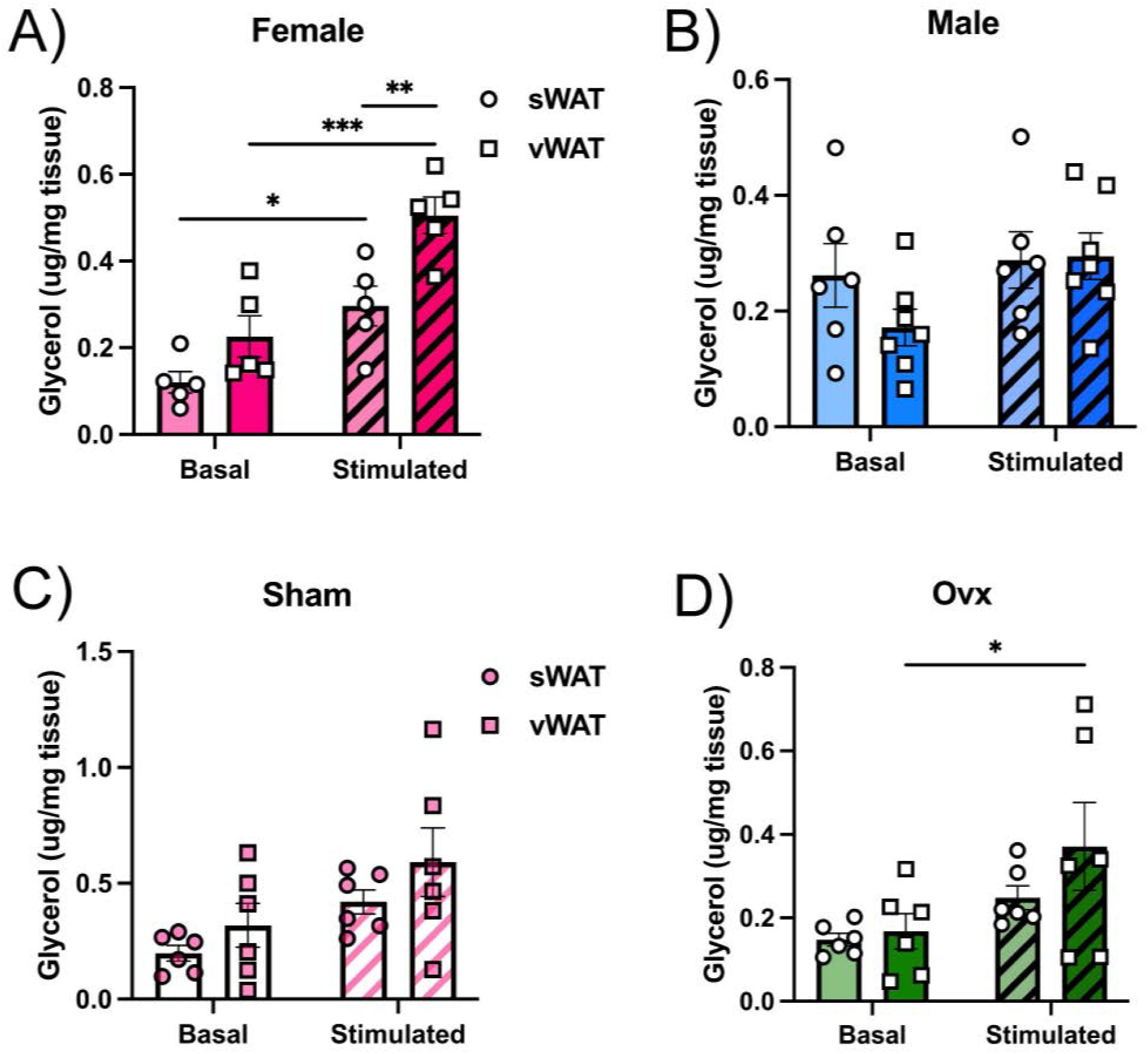
Depot comparison of isoproterenol-stimulated WAT lipolysis. Glycerol production at baseline and following 2h stimulation with isoproterenol in A) female; B) male; C) sham-operated; and D) ovx females. (n=5-7). *p<0.05, **p<0.01, ***p<0.001.

**Supp. Figure 4.**
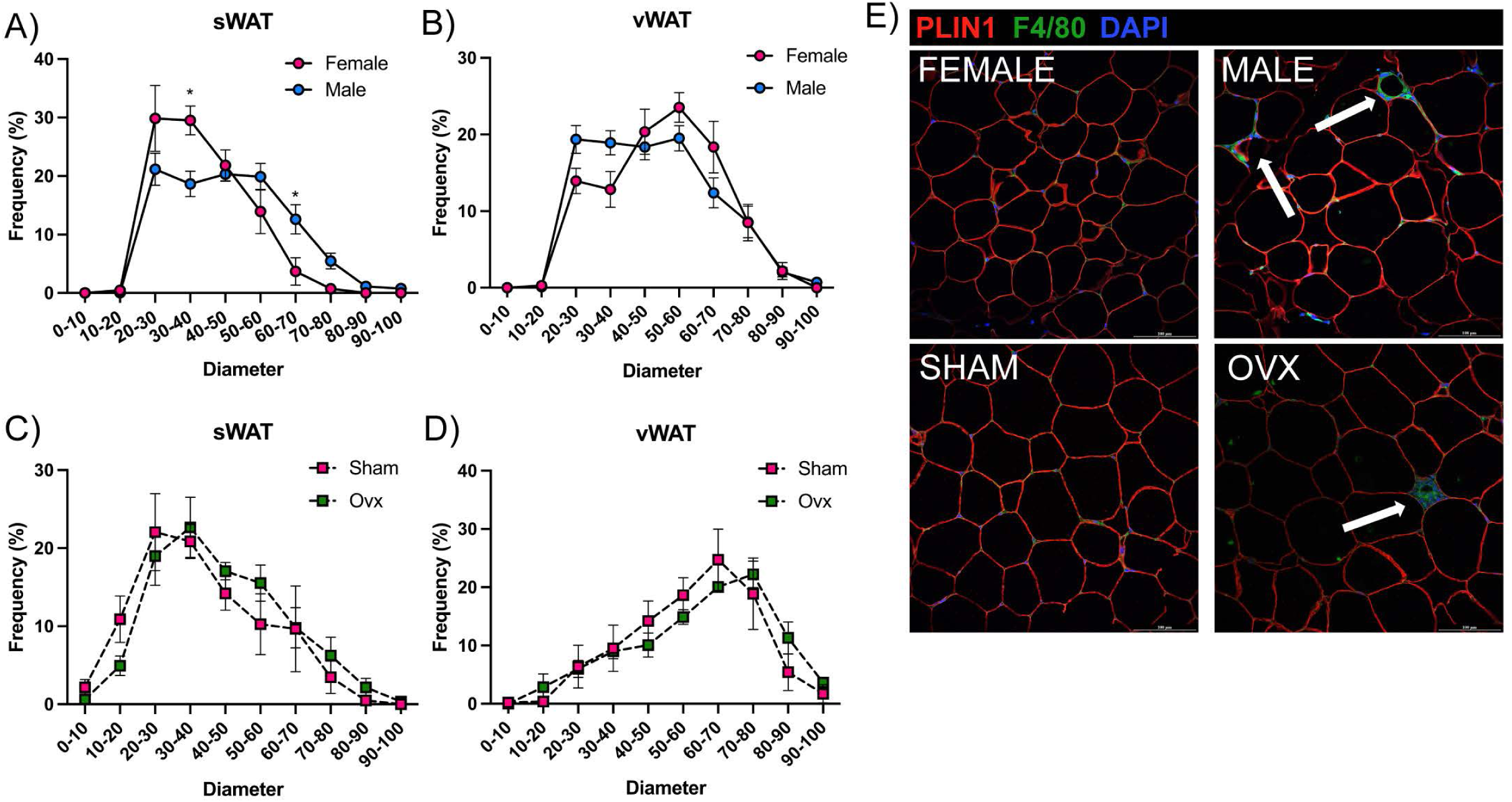
Sex differences in subcutaneous adipose size. A, C) Adipocyte size in sWAT of male, female, ovx, and sham mice, shown as frequency of adipocyte size. B, D) Adipocyte size in vWAT of male, female, ovx, and sham mice. E) Representative images of vWAT stained with perilipin 1 (membrane), F4/80 (macrophages) and Hoechst (nuclei) showing crown like structures in (arrows). (n=7-9). *p<0.05

## REFERENCES

1. Schnurr TM, Jakupović H, Carrasquilla GD, Ängquist L, Grarup N, Sørensen TIA, et al. Obesity, unfavourable lifestyle and genetic risk of type 2 diabetes: a case-cohort study. Diabetologia. 2020 Jul;63(7):1324–32.

2. Langenberg C, Sharp SJ, Franks PW, Scott RA, Deloukas P, Forouhi NG, et al. Gene-Lifestyle Interaction and Type 2 Diabetes: The EPIC InterAct Case-Cohort Study. Hattersley AT, editor. PLoS Med. 2014 May 20;11(5):e1001647.

3. Palmer BF, Clegg DJ. The sexual dimorphism of obesity. Mol Cell Endocrinol. 2015 Feb 15;0:113–9.

4. Steiner BM, Berry DC. The Regulation of Adipose Tissue Health by Estrogens. Front Endocrinol (Lausanne). 2022 May 26;13:889923.

5. Jeon J, Jung KJ, Jee SH. Waist circumference trajectories and risk of type 2 diabetes mellitus in Korean population: the Korean genome and epidemiology study (KoGES). BMC Public Health. 2019 Dec;19(1):741.

6. Greendale GA, Han W, Finkelstein JS, Burnett-Bowie SAM, Huang M, Martin D, et al. Changes in Regional Fat Distribution and Anthropometric Measures Across the Menopause Transition. The Journal of Clinical Endocrinology & Metabolism. 2021 Aug 18;106(9):2520–34.

7. Gurka MJ, Vishnu A, Santen RJ, DeBoer MD. Progression of Metabolic Syndrome Severity During the Menopausal Transition. JAHA [Internet]. 2016 [cited 2022 Aug 23];5(8). Available from: https://www.ahajournals.org/doi/epub/10.1161/JAHA.116.003609

8. Janssen I. Menopause and the Metabolic SyndromeThe Study of Women’s Health Across the Nation. Arch Intern Med. 2008 Jul 28;168(14):1568.

9. Purnell JQ, Urbanski HF, Kievit P, Roberts CT, Bethea CL. Estradiol Replacement Timing and Obesogenic Diet Effects on Body Composition and Metabolism in Postmenopausal Macaques. Endocrinology. 2019 Apr 1;160(4):899–914.

10. Lovre D, Lindsey SH, Mauvais-Jarvis F. Effect of menopausal hormone therapy on components of the metabolic syndrome. Ther Adv Cardiovasc Dis. 2017 Jan;11(1):33–43.

11. for the Women’s Health Initiative Investigators, Margolis KL, Bonds DE, Rodabough RJ, Tinker L, Phillips LS, et al. Effect of oestrogen plus progestin on the incidence of diabetes in postmenopausal women: results from the Women’s Health Initiative Hormone Trial. Diabetologia. 2004 Jul;47(7):1175–87.

12. Zucker I, Beery AK. Males still dominate animal studies. Nature. 2010 Jun;465(7299):690–690.

13. Virtue S, Vidal-Puig A. It’s Not How Fat You Are, It’s What You Do with It That Counts. PLoS Biol. 2008 Sep 23;6(9):e237.

14. Joe AWB, Yi L, Even Y, Vogl AW, Rossi FMV. Depot-Specific Differences in Adipogenic Progenitor Abundance and Proliferative Response to High-Fat Diet. Stem Cells. 2009 Oct;27(10):2563–70.

15. Jeffery E, Church CD, Holtrup B, Colman L, Rodeheffer MS. Rapid depot-specific activation of adipocyte precursor cells at the onset of obesity. Nat Cell Biol. 2015 Apr;17(4):376–85.

16. Wang QA, Tao C, Gupta RK, Scherer PE. Tracking adipogenesis during white adipose tissue development, expansion and regeneration. Nat Med. 2013 Oct;19(10):1338–44.

17. Jeffery E, Wing A, Holtrup B, Sebo Z, Kaplan JL, Saavedra-Peña R, et al. The Adipose Tissue Microenvironment Regulates Depot-Specific Adipogenesis in Obesity. Cell Metabolism. 2016 Jul;24(1):142–50.

18. Bilal M, Nawaz A, Kado T, Aslam MR, Igarashi Y, Nishimura A, et al. Fate of adipocyte progenitors during adipogenesis in mice fed a high-fat diet. Molecular Metabolism. 2021 Dec;54:101328.

19. Tang W, Zeve D, Seo J, Jo AY, Graff JM. Thiazolidinediones Regulate Adipose Lineage Dynamics. Cell Metabolism. 2011 Jul;14(1):116–22.

20. de Souza CJ, Eckhardt M, Gagen K, Dong M, Chen W, Laurent D, et al. Effects of Pioglitazone on Adipose Tissue Remodeling Within the Setting of Obesity and Insulin Resistance. Diabetes. 2001 Aug 1;50(8):1863–71.

21. Tang QQ, Lane MD. Adipogenesis: From Stem Cell to Adipocyte. Annu Rev Biochem. 2012 Jul 7;81(1):715–36.

22. Mikolajczak A, Sallam NA, Singh RD, Scheidl TB, Walsh EJ, Larion S, et al. Accelerated developmental adipogenesis programs adipose tissue dysfunction and cardiometabolic risk in offspring born to dams with metabolic dysfunction. American Journal of Physiology- Endocrinology and Metabolism. 2021 Nov 1;321(5):E581–91.

23. Kim SM, Lun M, Wang M, Senyo SE, Guillermier C, Patwari P, et al. Loss of White Adipose Hyperplastic Potential Is Associated with Enhanced Susceptibility to Insulin Resistance. Cell Metabolism. 2014 Dec;20(6):1049–58.

24. Park HT, Lee ES, Cheon YP, Lee DR, Yang KS, Kim YT, et al. The relationship between fat depot-specific preadipocyte differentiation and metabolic syndrome in obese women: Preadipocyte and metabolic syndrome. Clinical Endocrinology. 2012 Jan;76(1):59–66.

25. Nguyen HP, Lin F, Yi D, Xie Y, Dinh J, Xue P, et al. Aging-dependent regulatory cells emerge in subcutaneous fat to inhibit adipogenesis. Developmental Cell. 2021 May;56(10):1437–1451.e3.

26. Miyazaki Y, Mahankali A, Matsuda M, Mahankali S, Hardies J, Cusi K, et al. Effect of Pioglitazone on Abdominal Fat Distribution and Insulin Sensitivity in Type 2 Diabetic Patients. The Journal of Clinical Endocrinology & Metabolism. 2002;87(6):8.

27. Dennis JM, Henley WE, Weedon MN, Lonergan M, Rodgers LR, Jones AG, et al. Sex and BMI Alter the Benefits and Risks of Sulfonylureas and Thiazolidinediones in Type 2 Diabetes: A Framework for Evaluating Stratification Using Routine Clinical and Individual Trial Data. Diabetes Care. 2018 Sep 1;41(9):1844–53.

28. Nissen SE, Wolski K. Effect of Rosiglitazone on the Risk of Myocardial Infarction and Death from Cardiovascular Causes. N Engl J Med. 2007 Jun 14;356(24):2457–71.

29. Power ML, Schulkin J. Sex differences in fat storage, fat metabolism, and the health risks from obesity: possible evolutionary origins. Br J Nutr. 2008 May;99(5):931–40.

30. Holtrup B, Church CD, Berry R, Colman L, Jeffery E, Bober J, et al. Puberty is an important developmental period for the establishment of adipose tissue mass and metabolic homeostasis. Adipocyte. 2017 Jul 3;6(3):224–33.

31. Chau YY, Bandiera R, Serrels A, Martínez-Estrada OM, Qing W, Lee M, et al. Visceral and subcutaneous fat have different origins and evidence supports a mesothelial source. Nat Cell Biol. 2014 Apr;16(4):367–75.

32. do Carmo JM, da Silva AA, Moak SP, Browning JR, Dai X, Hall JE. Increased sleep time and reduced energy expenditure contribute to obesity after ovariectomy and a high fat diet. Life Sciences. 2018 Nov 1;212:119–28.

33. Giles ED, Jackman MR, Johnson GC, Schedin PJ, Houser JL, MacLean PS. Effect of the estrous cycle and surgical ovariectomy on energy balance, fuel utilization, and physical activity in lean and obese female rats. American Journal of Physiology-Regulatory, Integrative and Comparative Physiology. 2010 Dec;299(6):R1634–42.

34. Rogers NH, Perfield JW, Strissel KJ, Obin MS, Greenberg AS. Reduced Energy Expenditure and Increased Inflammation Are Early Events in the Development of Ovariectomy-Induced Obesity. Endocrinology. 2009 May 1;150(5):2161–8.

35. Cheng Y, Zhu H, Ren J, Wu HY, Yu JE, Jin LY, et al. Follicle-stimulating hormone orchestrates glucose-stimulated insulin secretion of pancreatic islets. Nat Commun. 2023 Nov 1;14(1):6991.

36. D’Eon TM, Souza SC, Aronovitz M, Obin MS, Fried SK, Greenberg AS. Estrogen Regulation of Adiposity and Fuel Partitioning. Journal of Biological Chemistry. 2005 Oct;280(43):35983–91.

37. Church CD, Berry R, Rodeheffer MS. Isolation and Study of Adipocyte Precursors. In: Methods in Enzymology [Internet]. Elsevier; 2014 [cited 2021 May 6]. p. 31–46. Available from: https://linkinghub.elsevier.com/retrieve/pii/B9780124116191000033

38. Rodeheffer MS, Birsoy K, Friedman JM. Identification of White Adipocyte Progenitor Cells In Vivo. Cell. 2008 Oct;135(2):240–9.

39. Søndergaard E, Espinosa De Ycaza AE, Morgan-Bathke M, Jensen MD. How to Measure Adipose Tissue Insulin Sensitivity. J Clin Endocrinol Metab. 2017 Jan 31;102(4):1193–9.

